# Non-destructive radiocarbon dating of bone

**DOI:** 10.1101/2025.03.26.645488

**Authors:** Katharina Luftensteiner, Laura van der Sluis, Maddalena Giannì, Peter Steier, Andrei Belinski, Romain Mensan, Maxim Kozlikin, Michael Shunkov, John Schulze, Katerina Douka, Thomas W. Stafford, Tom Higham

## Abstract

Since the 1950s, radiocarbon measurements have anchored archaeological chronologies dating back to 50,000 years, with bone collagen being a commonly dated material. Despite advances in collagen extraction protocols, the process remains destructive, requiring sampling by sawing, drilling or crushing of dry bone, that can damage or destroy physical features and archaeological evidence, and often consumes the entire specimen. While non-destructive approaches have recently been applied in ancient genomics and palaeoproteomics, no equivalent approach has been established for radiocarbon dating of bone. Here, we outline a non-destructive collagen extraction workflow that avoids invasive sampling (cutting or drilling) and produces no visible damage to the bone. By heating the bone in hot water for several hours, collagen is solubilized, and the resulting collagenous solution can be purified and AMS dated. Here we show that the amino acid composition, C/N atomic ratios, δ^13^C and δ^15^N values of the hot-water-extracted collagen are comparable to collagen isolated from the same bones using classic decalcification methodologies. The hot water extraction method was tested on various bones, ranging from the Bronze Age to the Middle and Upper Paleolithic periods, which had been dated previously using routine destructive methods that involved acid demineralization. Our results show that non-destructive collagen extraction, coupled with an additional purification step e.g., ultrafiltration, yields identical radiocarbon ages to those obtained via the routine destructive methods.

## Introduction

Bone is an attractive material for radiocarbon dating in archaeology and paleontology, because vertebrate remains often demonstrate human presence and activities, and are direct evidence of a site’s paleoecology, paleoenvironments, and extinction histories. Since the 1980s, radiocarbon (^14^C) dating methods for bone have steadily improved, as techniques were developed to standardize protocols^1^. The ∼20% organic component of bone predominately consists of collagen^1,2^, which is the preferred material for ^14^C dating. The inorganic, or mineral, fraction of bone is bioapatite (carbonate-hydroxy-calcium phosphate) that can readily exchange its carbonate fraction with environmental carbonates and produce inaccurate ^14^C measurements. In contrast, bone collagen amino acids do not exchange carbon with the environment, although its peptides can be contaminated by foreign organic compounds, e.g., humates bound to amino acids by the Maillard reaction^2^. Accurate ^14^C dating of bone collagen involves the removal of contaminants, e.g humic and fulvic acids, and the isolation of collagen-specific peptides, α-carboxyl carbon from primary amino acids or the secondary, imino acid, hydroxyproline. A routine method for ^14^C dating bone collagen usually comprises, (1) destructive sampling of the bone, tooth, antler or ivory through cutting or drilling; (2) demineralization in dilute HCl to remove hydroxyapatite and isolate bulk collagen, (3) extraction with NaOH or KOH to mobilize humic compounds, followed by an acid wash, 4) gelatinization in weakly acidic water to solubilize the collagen triple helix^3^ and finally, (5) lyophilization and combustion of the gelatin, graphitization of CO2, and ^14^C measurement. Throughout this paper we refer to this method as the “destructive method”. Additional methods designed to improve accuracy by removing contaminants or isolating a collagen-specific component include >30 kDa ultrafiltration of gelatin^4^, XAD-2 resin purification of hydrolyzed gelatin^5^, the chromatographic isolation of a specific amino acid, e.g., hydroxyproline^5,6^ and dating carboxyl carbon from the alpha-carboxyl group of primary amino acids^7,8^. The commonality of all these methods is that they start with destructive processing of bone.

Yet, archaeological and paleontological specimens are rare and finite. Given that most conventional sampling techniques are inherently destructive, better methods are needed to obtain biomolecular and ^14^C data without destroying these bones.

In recent years, in fields such as ancient genomics and palaeoproteomics (specifically ZooMS), efforts have been made to reduce the damage to artefacts and fossils by using non- or minimally-destructive techniques. Recently, minimally invasive techniques have been used to extract ancient DNA (aDNA) from bone without cutting or drilling bone^6^. Essel et al. (2024) extracted *in situ* aDNA from a 25,000-year-old deer tooth pendant from Denisova Cave without destructive sampling, and obtained the mitochondrial genome from both the human individual who probably wore the pendant, as well as the deer itself. Similarly, ZooMS studies have tested different non-destructive methods of extracting and isolating collagen peptides from Medieval parchments using rubber erasers^7^ or buffers,^8^ and collected peptides from inside plastic bags originally containing bones^12^. The use of polishing films^9^ has also proven successful in extracting enough proteins for proteomic analysis.

In radiocarbon dating, non-destructive approaches remain extremely rare. One of the few minimally-destructive methods that has been developed for radiocarbon dating uses plasma oxidation, which was first applied to rock art samples and later to mummified remains, organic objects and bone samples^10,11^. The method separates labile organic constituents from the substrate without visible physical damage, making the technique suitable for certain perishable artifacts and artwork. However, these methods have been primarily used for the direct extraction of carbon from lipids and cellulose, which lack the stable isotope and amino acid sequence data provided by proteins.

In 1991, Stafford^12^ published a paper about testing and applying new chemical methods for dating ancient bones, and included a small test on a method for isolating collagen using a hot water treatment. Based on the comparison of ^14^C measurements for single amino acids isolated from Upper Paleolithic bone using both demineralization and hot water extraction methods, they concluded that “…*non-destructive isolation of bone protein yielded ages concordant with dates on* [decalcified] *collagen*” and “…*well-preserved, irreplaceable bone artifacts or human skeletons may be datable by extracting protein by non-destructive methods*”.

Building on and expanding these early non-destructive attempts, we experimented with non-destructive extraction and dating of bone collagen by immersing whole bone fragments in hot water. We tested heating times and temperatures, and with using a subsequent purification step, e.g., ultrafiltration, prior to AMS dating. To characterize the extracted material and show that hot water extracts from whole bone have the amino acid composition of bone collagen, we applied a range of analytical methods. We show that the non-destructive protocol produces radiocarbon ages that are statistically identical to those obtained using the destructive alternative.

Detailed sample information and pretreatment steps are documented in the Supplementary Information (Table S1).

## Results

### Hot water extraction of the Hollis mammoth bone

Non-destructive, hot water extraction was tested initially on Hollis mammoth bone, which is used as a background in the Higham Radiocarbon Laboratory. The Hollis bone was recovered from permafrost in the Yukon territory, Canada and dates to the Middle Pleistocene, ca. 700 ka^13^. The bone is a source of chemically well-preserved collagen with no detectable ^14^C.

Ten small fragments of the Hollis mammoth bone weighing 429-to-547 mg were placed into 10 mL baked-out borosilicate tubes and covered in MilliQ™ water at pH 7. The tubes were heated on a hot plate at 75°C, for 1 to 10 hours. Each hour, a sub-sample was removed from the tube, and the water-soluble extract was frozen overnight at -20°C and then freeze-dried for 48 hours. The yields of the residue, which had the visual appearance of collagen (white, cotton-like), were determined through weighing using high precision balances. There was a positive correlation between heating time and “collagen” yield (Fig.1a). The maximum yield was reached after 10 hours of heating (sample R00001.69) and produced 25.56 mg of extracted “collagen”, a 5.28% weight yield that is approximately 50% of the average 10.27% weight collagen yield when Hollis bone is processed using routine Longin pretreatment chemistry following the destructive method.

**Figure 1.**
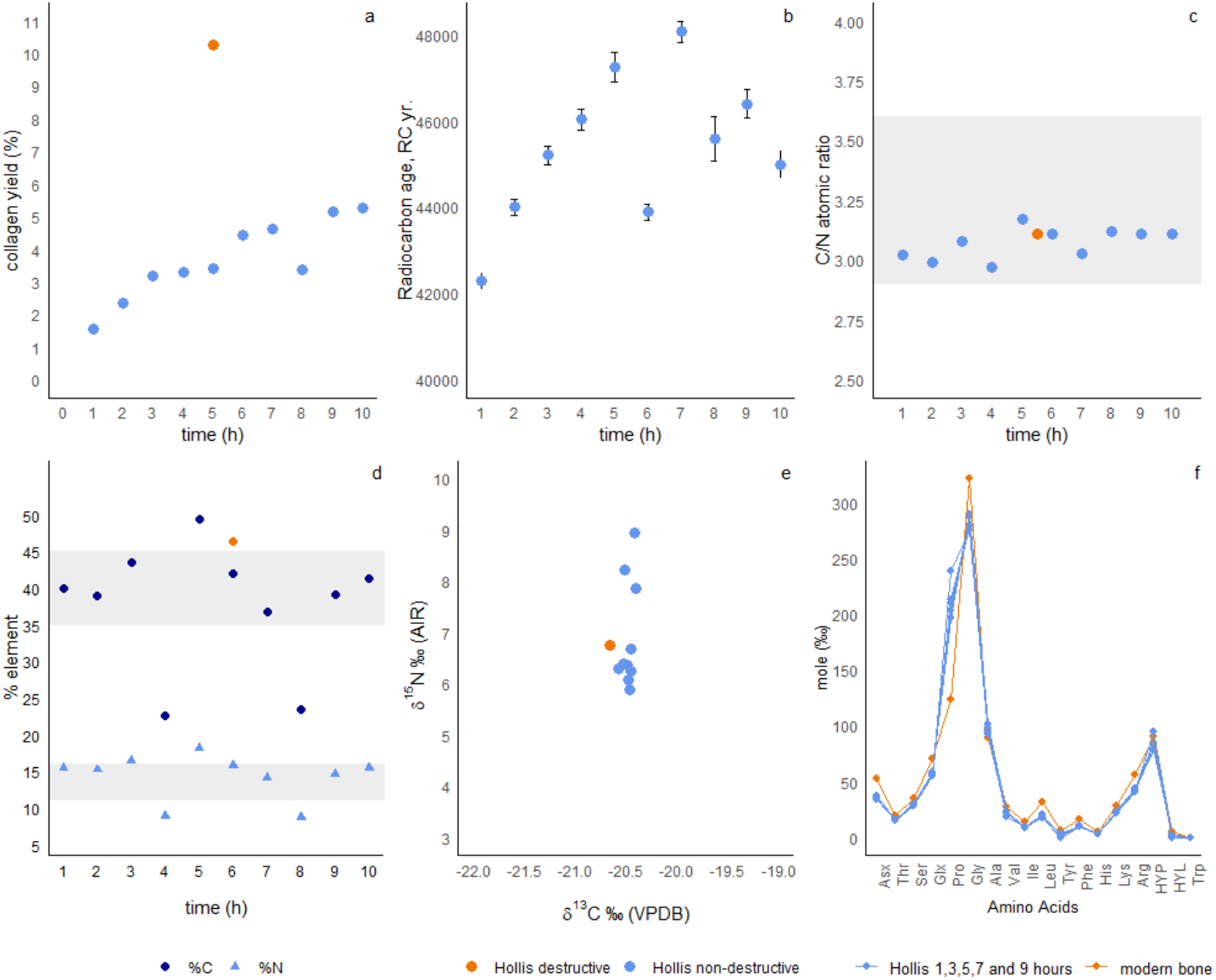
Analytical data for hot water extraction of 10 subsamples from the Hollis mammoth bone. Ten bone samples underwent hot water extraction for 1 to10 hours at 75°C. Comparative data generated using the destructive method are also shown. The graphs are; (a) collagen yield (% wt.); (b) radiocarbon ages, in years BP; (c) C/N_atomic_ values; (d) %N and %C vs. extraction time in relation to the time of treatment. Stable isotope data are shown (errors are ±0.2‰ for C and ±0.3‰ for N) (e) and amino acid compositions (f) are compared to modern human bone sample. Only collagen from 1, 3, 5, 7 and 9 hour extraction times were analyzed for their amino acid composition and are shown here.

To characterize the composition of the extracted material, we performed amino acid analyses, measured carbon and nitrogen stable isotope values, and C/N_atomic_ values on freeze-dried collagen. C/N_atomic_ values on bulk collagen or gelatin approximate protein quality. Acceptable C/N_atomic_ values outside the range of 2.9–3.6 may indicate the presence of non-collagenous proteins, degradation of the collagen molecule or both^14^. The theoretical, median C/N_atomic_ value is 3.243^15^. Bone collagen extraction of the Hollis mammoth bone using our laboratory’s destructive method (ABA with ultrafiltration) has C/N values averaging 3.12. C/N values for the 10 Hollis samples are 3.0–3.2 and consistent with bone collagen (Fig. 1c).

When we compare the non-destructively extracted collagen with the amino acid profile of modern bone, we observe similar values in amino acid composition, an indication that hot water extracts mirror the profile of Type 1 collagen (Fig. 1f). The δ^13^C and δ^15^N values from both methods are consistent with previously-obtained stable isotope data for the Hollis mammoth bone collagen. The Hollis samples extracted for 8, 9 and 10 hours show elevated δ^13^C and δ^15^N values. These stable isotope values may be higher due to non-collagenous leachates, a hypothesis tested by 30 kDa -filtering the 9-hour extract. The resulting δ^15^N values were 6.8‰, similar to those previously measured in our lab.

C/N_atomic_ values are indicative of intact collagen^16^. In well-preserved collagen, %C content ranges from 35–45%. Higher values indicate exogenous organic carbon. Degraded collagen shows carbon values under 30%. Nitrogen content in intact collagen should range between 11– 16%, which decreases proportionally with collagen degradation^17^. The Hollis standards showed very low values both in %C and %N following 4- and 8-hours hot water extractions (22.6 %C and 8.8 %N; 23.5 %C and 8.8 %N, respectively) although the remaining samples ranged from 14.1% to 18.2% in nitrogen content (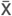 =15.72, *SD*=1.1) and from 36.7% to 49.4% in carbon content (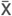 =41.36, *SD* = 3.41). While some samples lie outside the ranges suggested by van Klinken (1999)^17^, they are similar to Hollis samples processed using destructive methods in the Higham lab (%C 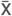 =46.43, *SD*=4.6)(Fig. 1d).

Overall, hot water extracts are collagenous; however, deviations from the norm may be due to unknown, non-collagenous organic matter dissolving in the solution.

To test for residual contamination, and without any further purification, we combusted each of the hot water-extracted samples on an EA (elemental analyser) and graphitized the generated CO2 using an AGE3 system at the Vienna Environmental Research Accelerator (VERA) facility, Faculty of Physics, University of Vienna. The graphites produced were measured on a National Electrostatics (NEC) 3-MV Pelletron tandem accelerator on two separate runs (see *Methods*) and the results are reported in Figure 1b and Table S3 in radiocarbon years BP.

The first Hollis ^14^C dates (samples 1-7) range from 42,282 ± 176 to 48,068 ± 249 BP (Fig. 1b). These contrast with ^14^C measurements on demineralized Hollis bone collagen that range between 50,031 ± 273 to 52,313 ± 377 BP, the laboratory’s background. Samples 8-10 ranged from 44,991 ± 308 to 46,398 ± 328 BP (Table S3). Therefore, we determine the amount of contamination to be 0.1–0.3% of modern carbon.

The differences between ^14^C ages obtained from simple hot water collagen extraction versus HCl demineralization were caused by the lack of ultrafiltration following the non-destructive protocol. The results from the hot water extraction were more than 40,000 BP, without any post-extraction cleaning of the bone collagen (i.e. no ultrafiltration or other purification steps), which are usually applied in radiocarbon dating of bone.

The initial results and the analytical data showed that hot water extraction alone yields an extract that is largely collagenous and with a yield that is up to -% of the demineralization process.

### Application to archaeological samples

The original results from the Hollis mammoth bone encouraged us to widen the study and apply the method to archaeological samples. We tested samples from two Bronze Age sites in the Russian north Caucasus (Ipatovo and Zaragizh) and two Palaeolithic sites, one in France (Abri Cellier) and one in the Russian Altai (Denisova Cave). Ipatovo and Abri Cellier bones yielded very low amounts of bone collagen and are not discussed further (see Supplementary Information, Table S2, Figure S1). Initial testing of samples from these sites, however, confirmed that for less well-preserved bones, collagen yields increased when the extraction temperature was increased from 75°C to 90°C (Table S2, Figure S1). This would, of course, be expected since reaction rates in chemistry double with every 10°C increase in temperature. We therefore modified our protocol and increased the hot water extraction temperature to 90°C. We compared these results with those obtained from the Destructive method to estimate the accuracy and reliability of ^14^C dates obtained using the non-destructive approach. We also added an ultrafiltration step to samples prepared using non-destructive collagen extraction to test the efficacy of further purifying hot water-extracted collagen. All samples were AMS ^14^C dated using the methods outlined above; analytical data and radiocarbon ages are presented in Table 2.

The bone samples that yielded enough collagen in the non-destructive extractions, yielded statistically identical ^14^C ages compared with the Destructive method after the collagen was purified by >30kDa ultrafiltration. Sample R00670 from Zaragizh was treated with four different extractions: hot-water extraction at 75°C and 90°C; hot water extraction at 90°C followed by ultrafiltration; and the Destructive method.

The ^14^C dates from the 75°C and 90°C hot water extracted fractions were 4513 ± 60 and 4799 ± 32 BP, respectively. Ultrafiltered collagen from the 90°C extraction was dated to 4906 ± 59 BP, which was statistically identical to the demineralized collagen ^14^C date (4928 ± 34 BP) (*T* = 0.1; df = 1, χ2 = 3.84).

These age estimates suggest that hot water extracted collagen must be further purified to obtain accurate ^14^C dates and remove contaminants, such as humates and fulvic acids. The Zaragizh tooth (R00668) was ^14^C dated using both demineralized and hot water extracted collagen fractions. The radiocarbon determinations were 5103 ± 32 and 5019 ± 32 BP, respectively, and statistically identical (*T* = 3.45; df 1, χ2=3.84)(Supplementary Information). This suggests that the sample was either contaminant-free or that the contaminant’s Fm (fraction modern) was identical to that of the dentine collagen we extracted.

We analyzed five samples from Denisova Cave, each of which was split into two subsamples; one subsample underwent hot water extraction, while the other was demineralized with HCl. Three samples had low water-extraction yields (Supplementary Information) and were not analyzed further. Two bones yielded higher amounts of collagen and were successfully ^14^C dated. The hot water-extracted fraction was ultrafiltered using >30 kDa ultrafilters and ^14^C dated. The ^14^C dates we obtained are close to the background limit for both hot water-extracted and demineralized collagen (Table 1). The analytical data for the two successfully dated samples from Denisova Cave (R00680 and R00683) are very similar across the different treatments and confirm the good chemical preservation of the two Denisova bones (Figure 2).

**Table 1:** Collagen masses (mg) of samples before and after >30 kDa ultrafiltration (UF). The “before” masses are milligrams of 90°C water extracted collagenous material before ultrafiltration. The “after” masses are milligrams after ultrafiltration.

| sample | site | Collagen yield-(mg) |  | %C |  | C/N |  |
| --- | --- | --- | --- | --- | --- | --- | --- |
|  |  | before UF | after UF | before UF | after UF | before UF | after UF |
| R00670 | Zaragizh | 9.32 | 2.13 | 53.28 | 42.9 | 3.36 | 3.1 |
| R00680 | Denisova Cave | 18.82 | 13.68 | 44.17 | 48 | 3.18 | 3.1 |
| R00683 | Denisova Cave | 3.04 | 2.04 | 48.97 | 42.4 | 3.2 | 3 |

**Figuer 2:**
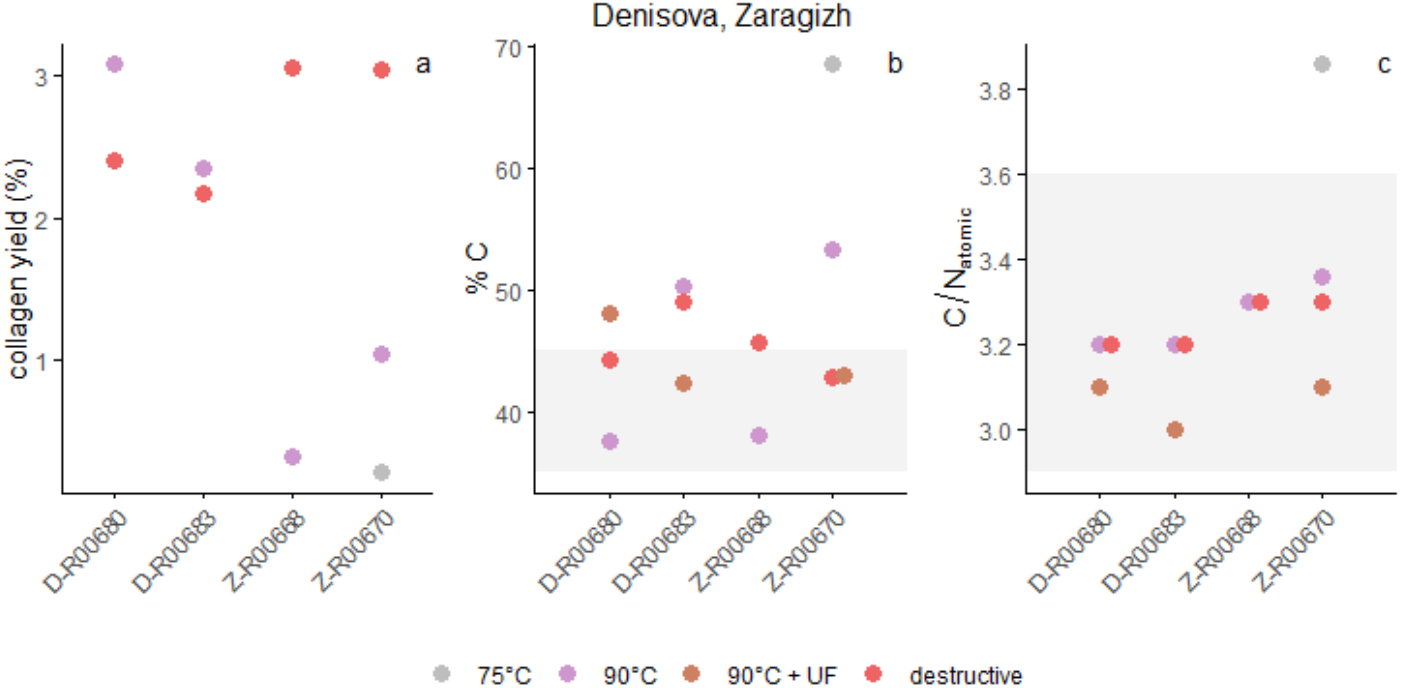
Collagen yield, %C and C/N ratios after hot water extraction. (a) Collagen yields after hot water extraction. Samples R00668 and R00670 underwent consecutive extraction rounds; the yields after each procedure are added to the total yield (Table S4); (b) carbon content (%C) of each sample; and (c) the C/N_atomic_ value. The grey area represents the broad range under which samples are considered well preserved^17^.

In summary, the archaeological dating results demonstrate that hot-water-extraction, followed by ultrafiltration using >30 kDa ultrafilters, enables accurate dating of fossil bones. Ultrafiltration, in any size range, reduces the bone’s collagen yield (Table 2) because smaller peptide fragments deriving from the bone collagen itself are eliminated. Future ^14^C dating will use XAD-2 resin purification of hydrolyzed gelatin, which will recover all amino acids and significantly increase yields (Stafford, et al 1988; 1991, van der Sluis et al. (2023).

**Table 2:** Radiocarbon ages and analytical data for bone samples dated by non-destructive and destructive (HCl demineralization) methods from Zaragizh and Denisova Cave. ND denotes the non-destructive protocol. ABA-UF – denotes the Destructive method with subsequent ultrafiltration. ND-UF denotes samples dated using the non-destructive collagen extraction protocol followed by ultrafiltration. Note that sample R00670 underwent consecutive hot water extraction with 75°C and 90°C. Collagen yield of the previous extraction round was added to the total yield (see Table S4).

|  | Zaragizh |  |  |  |  |  |  | Denisova |  |  |  |  |  |
| --- | --- | --- | --- | --- | --- | --- | --- | --- | --- | --- | --- | --- | --- |
| sample | R00670.1 | R00670.2 | R00670.3 | R00670.4 | R00668.1 | R00668.2 | R00668.3 | R00680.1 | R00680.2 | R00680.3 | R00683.1 | R00683.2 | R00683.3 |
| mass (mg) | 1126 | 1126 | 1126 | 1126 | 1229 | 1224 | 564 | 914 | 877 | 914 | 515 | 490 | 515 |
| Protocol | ND | ND | ABA-UF | ND-UF | ND | ND | ABA-UF | ND | ABA-UF | ND-UF | ND | ABA-UF | ND - UF |
| Time (h) | 9 | 9 | x | 9 | 9 | 9 | x | 9 | x | 9 | 9 | x | 9 |
| Temp. °C | 75 | 90 | x | 90 | 75 | 90 | x | 90 | x | 90 | 90 | x | 90 |
| Yield (mg) | 2.38 | 11.7 | 34.18 | 2.13 | 0.56 | 3.4 | 33.7 | 28.15 | 21.04 | 13.68 | 12.09 | 10.62 | 2.04 |
| Yield (% wt.) | 0.21 | 1.04 | 3.04 |  | 0.05 | 0.28 | 5.98 | 3.08 | 2.40 |  | 2.35 | 2.17 |  |
| d<sup>15</sup>N (‰) | 11.3 | 11.0 | 10.8 | 10.6 | x | 10.8 | 10.6 | 6.0 | 5.9 | 5.3 | 9.0 | 9.2 | 8.7 |
| d<sup>13</sup>C (‰) | -20.4 | -19.7 | -19.6 | -19.7 | x | -19.1 | -19.2 | -19.7 | -19.7 | -19.7 | -19.4 | -19.3 | -19.3 |
| % N | 20.7 | 18.5 | 15.1 | 16.2 | x | 13.5 | 16.4 | 13.6 | 16.2 | 17.8 | 18.3 | 17.8 | 16.5 |
| % C | 68.5 | 53.3 | 42.8 | 42.9 | x | 38.1 | 45.7 | 37.6 | 44.2 | 48.0 | 50.2 | 49.0 | 42.4 |
| C/N | 3.9 | 3.4 | 3.3 | 3.1 | x | 3.3 | 3.3 | 3.2 | 3.2 | 3.1 | 3.2 | 3.2 | 3.0 |
| VIE- number | 1158 | 1159 | 1160 | 1161 | x | 1156 | 1157 | 1165 | 1166 | 1167 | 1170 | 1171 | 1172 |
| <sup>14</sup>C age BP | 4513 | 4799 | 4928 | 4906 | x | 5103 | 5019 | >47700 | >48600 | >42500 | 45301 | >52100 | >38300 |
| ±1 se | 60 | 32 | 34 | 59 |  | 32 | 32 |  |  |  | 1471 |  |  |

### Changes in surface morphology

We also documented whether or not macroscopic changes were evident after the hot water extractions. We observed no visible morphological changes, only small variations in colour that were due mostly likely to the removal of trace sediments (Figure 4). Photographs of all of the samples are shown in Supplementary Figure S4.

**Figuer 3:**
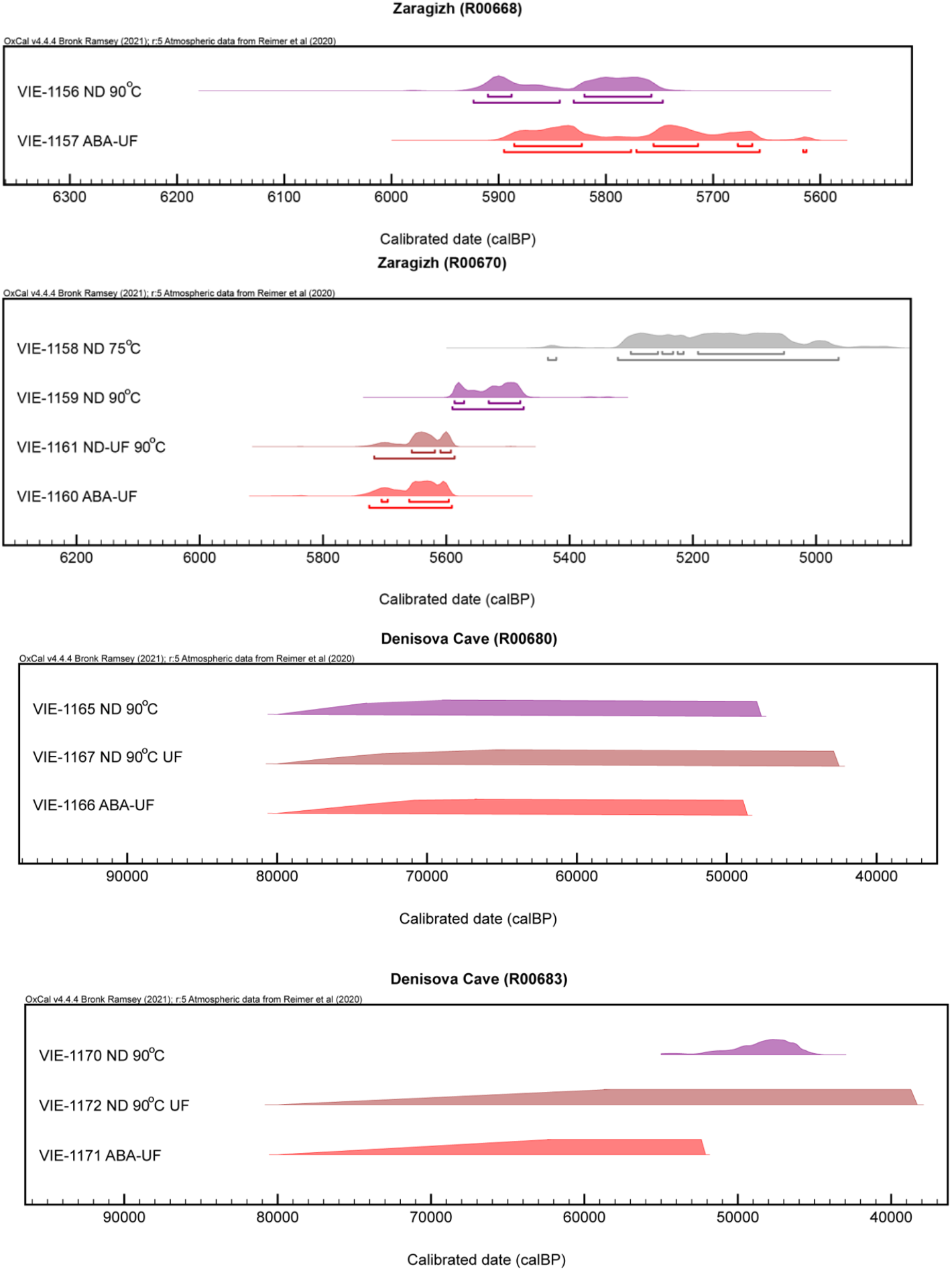
^14^C ages of archaeological samples calibrated as years before present (cal BP). ‘ND’ refers to the non-destructive hot water treatments, ‘ABA’ to the destructive protocol. ‘UF’ denotes samples that were ultrafiltered.

**Figure 4.**
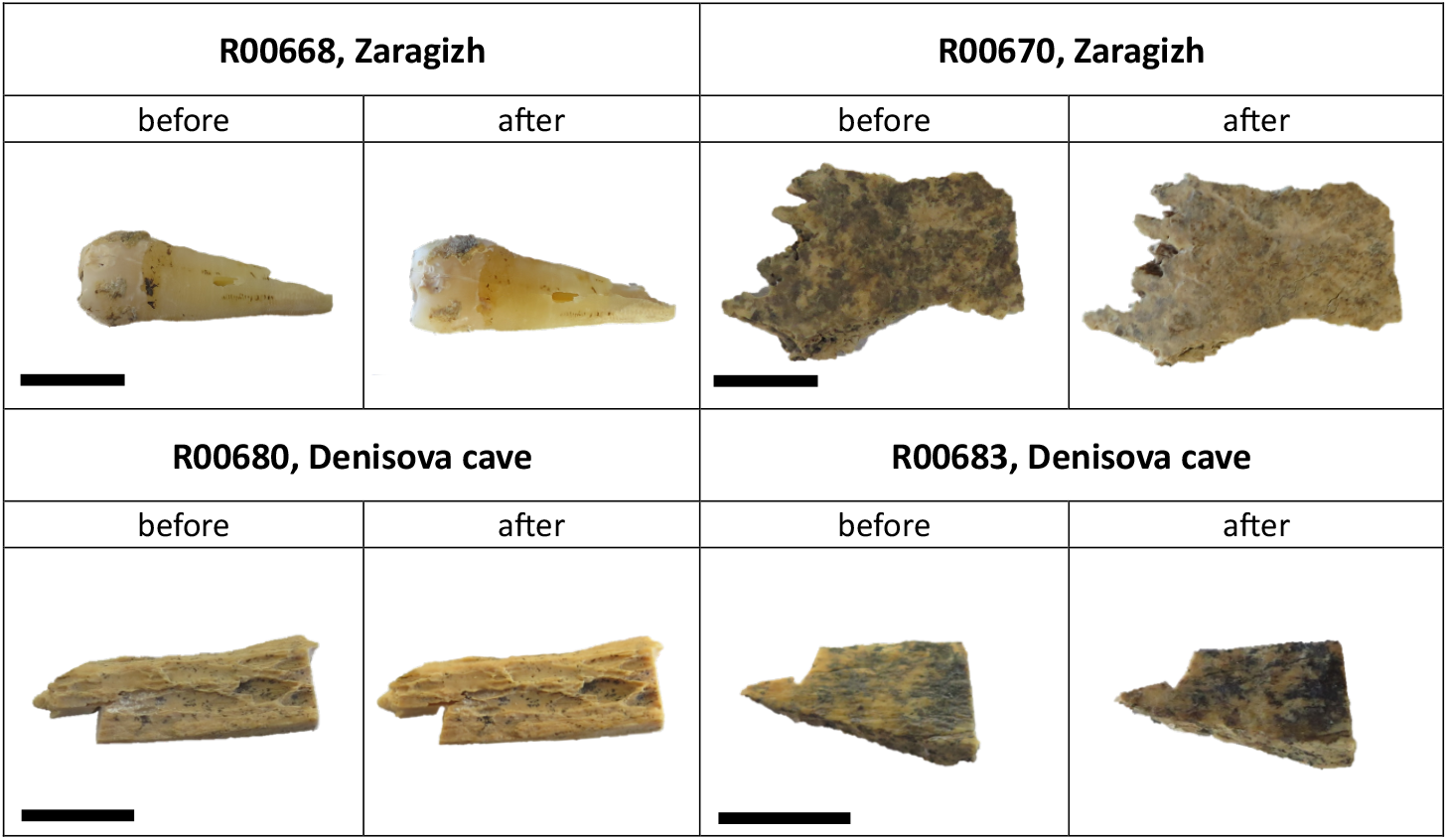
Photographs of samples taken before and after non-destructive hot water extraction. The black bar represents 1 cm. Note that colors are not directly comparable, as; (1) adhering sediment was washed off during the extraction process and; (2) the photographs were taken at slightly different angles, with different light settings and camera adjustments. Essentially no substantive differences in surface morphology can be seen. Note that for the tooth sample R00668 only the root was covered in MilliQ™ water, and the crown was only affected by evaporation.

## Discussion

In this study we have shown that collagen from bone and tooth dentine can be dated accurately by using a non-destructive, hot water extraction method. While accurate results may be obtained from hot-water-extracted collagen, further purification (ultrafiltration, treatment with XAD-resin, or extraction of hydroxyproline) should be performed regardless of how well preserved the collagen appears.

Previous studies have demonstrated that environmental temperature, soil pH and composition, water presence and microbial activity act together to affect collagen degradation in ancient bones^18,19^. In controlled experiments, collagen degradation has been simulated by heating bones to 90°C, comparable to our experimental procedure^20–22^. However, critical values for key parameters indicating collagen preservation quality were only reached after prolonged exposure, typically 12 to 14 days^21^. Since our samples were heated for ≤10 hours, we suggest that our treatment likely has no detrimental impact on collagen integrity.

Our experimental protocol parallels those on DNA degradation by heating, specifically the breaking of peptide bonds. DNA degradation occurs at lower temperatures when DNA is in solution rather than dry, generally at 100-110°C^23^. However, direct comparisons to aDNA are limited, because thermal degradation studies are made on modern DNA, rather than aDNA.

Non-destructive hot water extraction primarily releases the water-soluble fractions of bone collagen, while the destructive collagen extraction uses HCl to demineralize and recover bulk collagen.^3^ We observed no macroscopic damage, confirming that hot water extraction did not adversely affect sample surfaces, osteological landmarks or dimensions. However, we acknowledge that our study is limited by its focus on macroscopic evaluation, and further research is required to assess whether or not this extraction method impacts microscopic integrity, including the preservation of aDNA or other biomolecules. We suspect that the hot water extraction is likely to have a negligible effect given that Essel et al.^6^ observed that no aDNA was recoverable from artefacts heated to 90°C^24^.

In addition to the ethical component of the new approach that eliminates the destruction of unique remains, our protocol offers the advantage of a reduction in treatment time for preparing bone collagen samples, compared with destructive methods. Routine protocols of the Destructive method usually require multiple days. Drilling or crushing 18-24 bone samples, a typical grouping of samples that are treated together, often requires at least one full day, while decalcification can take up to three days or longer^25^. Gelatinization requires between 1-6 hours, but most laboratories undertake a longer gelatinization to ensure higher yields are achieved. In contrast, our method requires minimal setup and enables extraction within a 9-hour timeframe. By incorporating an ultrafiltration step immediately after extraction, all preparative steps prior to freeze-drying can be completed in 2 days, reducing the pretreatment protocol by ∼72-96 hours and making the approach time and cost efficient, while preserving the integrity of the sample.

## Conclusions

- Collagenous material can be extracted from archaeological and paleontological bones and teeth without visible, macroscopical damage by treating the bones in 90°C MilliQ™ water for several hours.
- Stable isotope values, nitrogen and carbon content and C/N values for the hot water extracts are comparable to those obtained through the Destructive method using HCl demineralization.
- Well-preserved bones and teeth across the radiocarbon age range can be dated using the non-destructive approach. Following hot-water extraction, the addition of ultrafiltration or XAD-2 purification, results to accurate radiocarbon ages.
- This non-destructive method will enable accurate dating of precious artefacts, ornaments and other archaeologically valuable objects or museum artefacts, as well as very small vertebrate fossils that were previously not dateable by destructive ^14^C methods.

## Supporting information

Supplementary Information - Non Destructive dating

## Acknowledgements

We thank the Faculty of Life Sciences, University of Vienna, and the office of the Rektor for their financial and logistical support, as well as all staff and students at the Douka-Higham laboratories at the University of Vienna. We acknowledge Dr. Biaslan Atabiev (Institute of the Archaeology of the Caucasus, Nalchik) for allowing us to analyse the material from Zaragizh.

## Methods overview

Two approaches were followed; 1) the non-destructive method using hot water to extract collagen from bones, and; 2) the Destructive method using HCl demineralization to isolate collagen from archeological bone and teeth.

### Non-destructive collagen extraction using hot-water extraction of whole bone

Samples with significant surface detritus were cleaned by air abrasion using aluminum oxide powder. Each sample was photographed before and after processing.

All borosilicate glassware was baked at 450°C for 6 hours. Ultrapure MilliQ™ water was used for all solutions. Vials and beakers were placed on a heating block with temperature and duration adjusted to either 75°C or 90°C for between 1 and 10 hours. To minimize evaporation, glassware was covered with glass lids, Petri dishes or aluminium foil (depending on the size of the glass tube/beaker). When necessary, ultrapure MilliQ™ water was added to adjust for water loss due to evaporation. After hot water extraction the bone samples were removed and air-dried under the fume hood overnight. The aqueous supernatant was frozen (−20°C) and then freeze dried.

### Destructive (routine) collagen extraction using HCl demineralization

The customary radiocarbon dating protocol of bone at the University of Vienna laboratories follows the procedure outlined by Brock et al. (2010)^25^. For all chemical pretreatments, glass beakers and centrifuge tubes were also baked at 450°C for 6 hours. We recorded all weights and yields of collagen for each step of the treatment process (see Table S1 and S2).

Bones for destructive radiocarbon dating were cleaned by air abrasion with aluminum oxide powder and then crushed. In this project, the size of archaeological samples varied and sizes exceeded the values in destructive protocols to avoid altering sample morphology and reproduce the sample sizes encountered with actual fossils (490 – 1126 mg). No organic solvent extractions were applied because the bone specimens had not been treated with consolidants or glues.

Samples were initially treated with 0.5M HCl for 12 –24 hours, during which the acid was changed several times. Depending on the bone’s preservation and size, demineralization was done overnight either at room temperature or at 4°C to slow down the reaction process (see supplements). After rinsing with MilliQ™ water, the samples were treated with a 0.1M NaOH solution for 30 minutes at room temperature, followed by three rinses with MilliQ™ water. The alkali step removes a significant amount of humates. The final step was washing the demineralized collagen with 0.5M HCl for 15 minutes at room temperature. After final rinsing with MilliQ™ water, a pH3 (HCl) MilliQ™ solution was added and the samples were heated at 75°C for 20 hours. This gelatinization step solubilizes collagen, after which the gelatin solution was filtered through a 45-90 µm Ezee™ filter made by Elkay (UK). The filtrate was directly transferred into a pre-cleaned >30 kDa Sartorius ultrafilter, after Brock et al. (2007)^26^ and centrifuged at 2700 rpm until approximately 1mL of the fraction remained. Ezee filters and ultrafilters were cleaned as described in Brock et al. (2007)^26^. Through this process the high molecular weight components (>30 kDa) were separated from the low MW fractions (<30 kDa, which is mostly degraded collagen, salts and contaminants. The >30 kDa fraction was freeze dried and retained for AMS dating and yielded a similar appearance to what we routinely observe, i.e. fluffy and white.

