## Supplementary Information - Non Destructive dating for "Non-destructive radiocarbon dating of bone"

1. **Materials**

The bones and one tooth used in this study come from multiple archaeological sites and range in age from the Palaeolithic to the Bronze Age. These specimens include:

- Hollis mammoth bone, Middle Pleistocene (*n*=10)
- VIRI I whale bone, consensus age 8331 ± 6 BP[1] (*n*=3)
- Ipatovo 3 and Zaragizh, north Caucasus, Russia, Bronze Age (*n*=4)
- Abri Cellier, France, Upper Palaeolithic (*n*=3)
- Denisova Cave, Russia, Middle and Upper Palaeolithic (*n*=7)

Sample information, masses, chemistry used and $^{14}C$ and analytical measurements are presented in Table S1.

### 1.1. Internal Laboratory Standards

#### 1.1.1 Hollis mammoth bone

The Hollis mammoth is used as a radiocarbon background bone in the Higham Laboratory, and is measured regularly to quantify the background value for bones and teeth dated. The Hollis bone was used to test the protocol for non-destructive collagen extraction. Hollis is a *Mammuthus primigenius* pelvis discovered in a placer gold mine in the Klondike region of central Yukon, Canada[1] and has been dated to the Middle Pleistocene, approximately 700,000 years BP, using fission-track dating of two tephra layers at the site[2] and an early Middle Pleistocene tephra correlated to the locality[3]. AMS $^{14}C$ dating of Hollis bone collagen verified its age is > 50,000 years and is below $^{14}C$ detection limits. The Hollis mammoth bone has excellent chemical and physical preservation and minimal collagen degradation due to its permafrost burial. Published collagen yields are 7.9% wt.[1]; however, our yields for ultrafiltration are higher. Additionally, the Hollis bone requires minimal to no physical cleaning due to its pristine condition.

#### 1.1.2 VIRI I Whale Bone

[1] All determinations throughout this paper given with 'BP' are conventional radiocarbon determinations, expressed in BP after Ref 12.

The second laboratory standard is VIRI I bone from the skull of an unidentified whale collected in 1997 from Svalbard Island, Norway. "VIRI" refers to the Fifth International Radiocarbon Intercomparison, an interlaboratory collaboration designed to assess and compare the accuracy and precision of radiocarbon dates among labs worldwide. The VIRI-I whale has a consensus age of 8331 ± 6 radiocarbon years BP[3].

#### 1.2. Archaeological samples

##### 1.2.1. Ipatovo 3 and Zaragizh, North Caucasus, Russia

We obtained Bronze Age bones from the sites of Ipatovo 3 and Zaragizh, North Caucasus, Russia. Ipatovo is a small town approximately 120 kilometers northeast of Stravropol in the North Caucasus, in the northern steppe region. Near this town is a large burial mound or "Kurgan" that was excavated during rescue excavations between 1998 and 1999. The site is located within middle steppe and dry steppe vegetation zones[5]

##### 1.2.2 Abri Cellier, France; Denisova Cave, Russia

Numerous Palaeolithic sites in the Dordogne region of southwest France are concentrated along the Vézère river. They have yielded extensive evidence of Upper Palaeolithic human occupation, and important remains of figurative art and symbolic artefacts[4]. Among notable sites yielding important cave art is the large rock shelter Abri Cellier in Tursac, Dordogne. Despite uncertainties regarding its chronology, Abri Cellier has long been recognized as one of the key sites containing early art from the European Aurignacian[5]. Abri Cellier is a collapsed rock shelter located on the promontory separating the Vimont valley and the Combe de Vergne in the commune of Tursac. Facing south-east, the shelter is located on a rocky terrace overlooking the right bank of the Vézère valley. Following an initial operation by Denis Peyrony in 19056, it was during the summer of 1927 that systematic excavations took place under the direction of George Collie for Beloit College in the USA[7]. In 2014, the late Prof. Randall White re-opened the site aiming at evaluating what, if any, archaeological potential remained. White and his team, undertook initially cleaning of the site down to bedrock and exposed sagittal stratigraphic sections remaining at both the eastern and western extremities of the shelter. Finally, the eastern extremity of the shelter yielded a small zone of intact stratified deposits, which they excavated over a total area of about 2m$^2$. The eastern profile revealed three archaeological layers, separated from each other by sterile layers related to roof collapses. At the base of the sequence, right on the bedrock, discovery of an *in situ* modified block suggested strong affinities of the layer with the Early Aurignacian. The second archaeological level was characterized by its black color (US 102) and was topped by a brown level (US 100), containing archaeological material associated with very large collapsed blocks that form the morphology of the present slope. The stratigraphy is coherent with the results from the old excavations in which the richest layer, sitting directly on bedrock, yielded an Early Aurignacian with Aurignacian blades and split-base points and modified blocks. The Abri Cellier samples in this paper originate in White´s excavations.

Denisova Cave, located in the Siberian Altai Mountains of Russia, is a key site understanding the relationship of human groups that lived in Eurasia during the Middle and Late Pleistocene[8]. DNA sequencing of human remains discovered at the site has identified a previously unknown hominin group, the Denisovans[8,9]. Additionally, high-coverage genomes from both Neanderthal and Denisovan fossils show evidence of interbreeding between these hominins[10].

Denisova Cave consists of three chambers (Main, East, and South) that vary in sedimentary composition and have independent stratigraphic numbering systems. Most directly dated human remains from the site yielded infinite (>50 ka BP) radiocarbon ages, indicating they are associated with strata beyond the limit of $^{14}C$. A horse tooth from layer 9.2, and a human bone labelled “Denisova 14” from layer 9.3 in the East Chamber, gave finite radiocarbon ages of 45500 ± 2300 and 46300 ± 2600 BP[11]. The samples used in this study are from the upper layers of the East Chamber (layers 9.1-9.3) and were excavated between 2004 and 2015.

**Table S1 :** Summary of samples used in this study. The protocols used are coded ND for the non-destructive hot water extraction approach and ABA for the conventional destructive treatment that uses HCl demineralization to isolate collagen. UF indicates an ultrafiltered sample. A blank space indicates that no purification step was added. If material was taken for stable isotopes (SI), radiocarbon dating ($^{14}C$) or amino acid analysis (AAA). measurements are noted by a ✓ for performed or X indicating not analyzed.

| R number | Site | Context | Cultural association | Sample information | Mass (mg) | Protocol used | extraction time (h) | Water temp (°C) | SI | $^{14}C$ | AAA |
|---|---|---|---|---|---|---|---|---|---|---|---|
| **R00001.47** | Hollis | Standard | | *Mammuthus primigenius* pelvis | 440 | ND | 1 | 75 | ✓ | ✓ | ✓ |
| **R00001.48** | Hollis | Standard | | " | 513 | ND | 2 | 75 | ✓ | ✓ | x |
| **R00001.49** | Hollis | Standard | | " | 504 | ND | 3 | 75 | ✓ | ✓ | ✓ |
| **R00001.50** | Hollis | Standard | | " | 547 | ND | 4 | 75 | ✓ | ✓ | x |
| **R00001.51** | Hollis | Standard | | " | 483 | ND | 5 | 75 | ✓ | ✓ | ✓ |
| **R00001.52** | Hollis | Standard | | " | 429 | ND | 6 | 75 | ✓ | ✓ | x |
| **R00001.53** | Hollis | Standard | | " | 523 | ND | 7 | 75 | ✓ | ✓ | ✓ |
| **R00001.67** | Hollis | Standard | | " | 514 | ND | 8 | 75 | ✓ | ✓ | x |
| **R00001.68** | Hollis | Standard | | " | 518 | ND | 9 | 75 | ✓ | ✓ | ✓ |
| **R00001.69** | Hollis | Standard | | " | 484 | ND | 10 | 75 | ✓ | ✓ | x |
| **R00001.68.1** | Hollis | Standard | | " | 480 | ND-UF | 9 | 90 | ✓ | ✓ | ✓ |
| **R00040.21** | VIRI | Standard | | Whale cranium | 465 | ND | 2 | 75 | ✓ | x | x |

| R number | Site | Context | Cultural association | Sample information | Mass (mg) | Protocol used | extraction time (h) | Water temp (°C) | SI | $^{14}$C | AAA |
|---|---|---|---|---|---|---|---|---|---|---|---|
| **R00040.22** | VIRI | Standard | | " | 498 | ND | 5 | 75 | ✓ | x | x |
| **R00040.23** | VIRI | Standard | | " | 498 | ND | 7 | 75 | ✓ | x | x |
| **R00668.1** | K1-B2-PM2.1 | Kurgan 1 Burial 2 | Bronze Age | *Homo sapiens* tooth | 1229 | ND | 9 | 75 | x | x | x |
| **R00668.2** | K1-B2-PM2.2 | " | " | " | 1224 | ND | 9 | 90 | ✓ | ✓ | x |
| **R00668.3** | K1-B2-PM2.3 | " | " | " | 564 | ABA-UF | x | x | ✓ | ✓ | x |
| **R00669.1** | K1-B2-cf.1 | " | " | " | 1077 | ND | 9 | 75 | x | x | x |
| **R00669.2** | K1-B2-cf.2 | " | " | " | 1051 | ND | 9 | 90 | ✓ | x | x |
| **R00669.3** | K1-B2-cf.3 | " | " | " | 1046 | ABA-UF | x | x | ✓ | x | x |
| **R00670.1** | K1-B1-cf.1 | Kurgan 1 Burial 1 | " | " | 1126 | ND | 9 | 75 | ✓ | ✓ | x |
| **R00670.2** | K1-B1-cf.2 | " | " | " | 1072 | ND | 9 | 90 | ✓ | ✓ | x |
| **R00670.3** | K1-B1-cf.3 | " | " | " | 1036 | ABA-UF | x | x | ✓ | ✓ | x |
| **R00670.4** | K1-B1-cf.3 | " | " | " | 1072 | ND-UF | 9 | 90 | ✓ | ✓ | x |
| **R00667.1** | K5-4/6.1 | Kurgan 5 4/6 | Maikop | *Homo sapiens* bone | 191 | ND | 7 | 75 | x | x | ✓ |
| **R00667.2** | K5-4/6.2 | " | " | " | 187 | ND | 7 | 90 | ✓ | x | x |

| R number | Site | Context | Cultural association | Sample information | Mass (mg) | Protocol used | extraction time (h) | Water temp (°C) | SI | $^{14}C$ | AAA |
|---|---|---|---|---|---|---|---|---|---|---|---|
| **R00667.3** | K5-4/6.3 | " | " | " | 188 | ABA | x | x | ✓ | x | x |
| **R00671.1** | P38830.1 | U28C ID2 OS Feuille 0 Layer trace 1 | Upper Palaeolithic | | 137 | ND | 9 | 75 | x | x | x |
| **R00671.2** | P38830.2 | " | " | | 136 | ND | 9 | 90 | x | x | ✓ |
| **R00671.3** | P38830.3 | " | " | | 133 | ABA | x | x | x | x | x |
| **R00672.1** | P38831.1 | " | " | | 351 | ND | 7 | 75 | x | x | x |
| **R00672.2** | P38831.2 | " | " | | 346 | ND | 7 | 90 | ✓ | x | ✓ |
| **R00672.3** | P38831.3 | " | " | | 342 | ABA-UF | x | x | ✓ | x | x |
| **R00673.1** | P38833.1 | " | " | | 1536 | ND | 7 | 75 | x | ✓ | x |
| **R00673.2** | P38833.2 | " | " | | 1487 | ND | 7 | 90 | ✓ | ✓ | x |
| **R00673.3** | P38833.3 | " | " | | 1391 | ABA-UF | x | x | ✓ | ✓ | x |
| **R00677.1** | DC03849.1 | Denisova Cave, East chamber Layer 9.2 | Upper Palaeolithic | Equidae | 2388 | ND | 9 | 90 | ✓ | x | x |
| **R00677.2** | DC03849.2 | " | " | " | 2319 | ABA-UF | x | x | x | x | x |
| **R00678.1** | DC03853.1 | Denisova Cave, East | " | " | 1712 | ND | 9 | 90 | ✓ | x | x |

| R number | Site | Context | Cultural association | Sample information | Mass (mg) | Protocol used | extraction time (h) | Water temp (°C) | SI | $^{14}$C | AAA |
|---|---|---|---|---|---|---|---|---|---|---|---|
| | | chamber Layer 9.2 | | | | | | | | | |
| **R00678.2** | DC03853.2 | " | " | " | 1643 | ABA-UF | x | x | x | x | x |
| **R00679.1** | DC03862.1 | Denisova Cave, East chamber Layer 9.2 | " | *Ovis* | 667 | ND | 9 | 90 | ✓ | x | x |
| **R00679.2** | DC03862.2 | " | " | " | 628 | ABA | x | x | ✓ | x | x |
| **R00680.1** | DC03875.1 | Denisova Cave, East chamber Layer 9.2 | " | Bison/Yak | 914 | ND | 9 | 90 | ✓ | ✓ | ✓ |
| **R00680.2** | DC03875.2 | " | " | " | 877 | ABA-UF | x | x | ✓ | ✓ | ✓ |
| **R00680.3** | DC03875.3 | " | " | " | 914 | ND-UF | 9 | 90 | ✓ | ✓ | ✓ |
| **R00681.1** | DC07878.1 | Denisova Cave, East chamber Layer 9.3 | " | *Crocuta*/*Panthera* | 509 | ND | 9 | 90 | ✓ | ✓ | x |
| **R00681.2** | DC07878.2 | " | " | " | 497 | ABA-UF | x | x | ✓ | ✓ | x |
| **R00682.1** | DC07897.1 | Denisova Cave, East chamber Layer | " | *Coelodonta antiquitatus* | 466 | ND | 9 | 90 | ✓ | x | x |

| R number | Site | Context | Cultural association | Sample information | Mass (mg) | Protocol used | extraction time (h) | Water temp (°C) | SI | $^{14}C$ | AAA |
|---|---|---|---|---|---|---|---|---|---|---|---|
| | | 9.3 | | | | | | | | | |
| **R00682.2** | DC07897.2 | " | " | " | 446 | ABA | x | x | x | x | x |
| **R00683.1** | DC07917.1 | Denisova Cave, East chamber Layer 9.3 | " | *Bos/Bison* | 515 | ND | 9 | 90 | ✓ | ✓ | ✓ |
| **R00683.2** | DC07917.2 | " | " | " | 490 | ABA-UF | x | x | ✓ | ✓ | ✓ |
| **R00683.3** | DC07917.3 | " | " | " | 515 | ND-UF | 9 | 90 | ✓ | ✓ | ✓ |

**Table S2. Laboratory data and results on samples processed in this project**. Sample weights, information on treatments, analytical data, and radiocarbon ages (RC years BP) are listed. P code denotes the pretreatment method. The protocols are coded ND for non-destructive hot water extraction and ABA is the Destructive method. UF denotes ultrafiltration. When information is missing, the sample yielded insufficient mass or poor quality of collagen and further analysis was impossible. "Collagen extracted" is the mass of protein isolated. "Collagen yield" is percent of bone that is collagen after chemical pre-treatment. Measurement precisions are ±0.3‰ for $\delta^{15}N$ and ±0.2‰ for $\delta^{13}C$. %N and %C = nitrogen and carbon yield on combustion. C/N is the atomic ratio of carbon to nitrogen. Radiocarbon dates are reported following the conventions of Stuiver and Polach (1977)[12].

| Lab number | Site | Mass (mg) | P code | Extraction time (h) | Water temp °C | Collagen yield (mg) | % collagen yield | $\delta^{15}N$ (‰) | $\delta^{13}C$ (‰) | % N | % C | C:N atomic ratio | $^{14}C$ age (BP) | ± value (1 σ) |
|---|---|---|---|---|---|---|---|---|---|---|---|---|---|---|
| **R00001.47** | Hollis | 440 | ND | 1 | 75 | 6.8 | 1.55 | 6.1 | -20.5 | 15.5 | 40.0 | 3.02 | 42282 | 176 |
| **R00001.48** | Hollis | 513 | ND | 2 | 75 | 12.02 | 2.34 | 6.3 | -20.6 | 15.2 | 39.1 | 2.99 | 44006 | 183 |
| **R00001.49** | Hollis | 504 | ND | 3 | 75 | 16.15 | 3.20 | 6.4 | -20.5 | 16.5 | 43.6 | 3.08 | 45204 | 213 |
| **R00001.50** | Hollis | 547 | ND | 4 | 75 | 18.12 | 3.31 | 6.7 | -20.5 | 8.9 | 22.6 | 2.97 | 46034 | 241 |
| **R00001.51** | Hollis | 483 | ND | 5 | 75 | 16.46 | 3.41 | 5.9 | -20.5 | 18.2 | 49.4 | 3.17 | 47250 | 332 |
| **R00001.52** | Hollis | 429 | ND | 6 | 75 | 19.02 | 4.43 | 6.2 | -20.5 | 15.8 | 42.1 | 3.11 | 43895 | 188 |
| **R00001.53** | Hollis | 523 | ND | 7 | 75 | 24.11 | 4.61 | 6.4 | -20.5 | 14.1 | 36.7 | 3.03 | 48068 | 249 |
| **R00001.67** | Hollis | 514 | ND | 8 | 75 | 17.44 | 3.39 | 8.9 | -20.4 | 8.8 | 23.5 | 3.12 | 45584 | 517 |
| **R00001.68** | Hollis | 518 | ND | 9 | 75 | 26.65 | 5.14 | 8.2 | -20.5 | 14.7 | 39.1 | 3.11 | 46398 | 328 |
| **R00001.68.1** | Hollis | 480 | ND-UF | 9 | 75 | 12.25 | 2.55 | 6.3 | -20.6 | 16 | 41 | 3 | 39858 | 266 |
| **R00001.69** | Hollis | 484 | ND | 10 | 75 | 25.56 | 5.28 | 7.9 | -20.4 | 15.5 | 41.4 | 3.11 | 44991 | 308 |
| **R00040.21** | VIRI I | 465 | ND | 2 | 75 | 6.3 | 1.35 | 17.2 | -18.6 | 5.1 | 15.3 | 3.51 | - | - |
| **R00040.22** | VIRI I | 498 | ND | 5 | 75 | 16.02 | 3.22 | 14.4 | -20.9 | 3 | 10.8 | 4.24 | - | - |

| Lab number | Site | Mass (mg) | P code | Extraction time (h) | Water temp °C | Collagen yield (mg) | % collagen yield | $\delta^{15}N$ (‰) | $\delta^{13}C$ (‰) | % N | % C | C:N atomic ratio | $^{14}C$ age (BP) | ± value (1 σ) |
|---|---|---|---|---|---|---|---|---|---|---|---|---|---|---|
| **R00040.23** | VIRI I | 498 | ND | 7 | 75 | 4.09 | 0.82 | 16.1 | -20 | 3.1 | 10.9 | 4.12 | - | - |
| **R00667.1** | Zaragizh | 191 | ND | 7 | 75 | 0.64 | 0.34 | - | - | - | - | - | - | - |
| **R00667.2** | Zaragizh | 187 | ND | 7 | 90 | 0.34 | 0.18 | 12.4 | -18.6 | 2.5 | 10.6 | 5 | - | - |
| **R00667.3** | Zaragizh | 188 | ABA | x | x | 2.77 | 1.47 | 13.8 | -17.3 | 13.6 | 39.3 | 3.37 | - | - |
| **R00668.1** | Zaragizh | 1229 | ND | 9 | 75 | 0.56 | 0.05 | - | - | - | - | - | - | - |
| **R00668.2** | Zaragizh | 1224 | ND | 9 | 90 | 3.4 | 0.28 | 10.8 | -19.1 | 13.5 | 38.1 | 3.29 | 5103 | 32 |
| **R00668.3** | Zaragizh | 564 | ABA-UF | x | x | 33.7 | 5.98 | 10.6 | -19.2 | 16.4 | 45.7 | 3.26 | 5019 | 32 |
| **R00669.1** | Zaragizh | 1077 | ND | 9 | 75 | 0.08 | 0.01 | - | - | - | - | - | - | - |
| **R00669.2** | Zaragizh | 1051 | ND | 9 | 90 | 1.79 | 0.17 | 11.2 | -20.1 | 8.0 | 26.3 | 3.85 | - | - |
| **R00669.3** | Zaragizh | 1046 | ABA-UF | x | x | 17.28 | 1.65 | 11.2 | -19.2 | 14.3 | 41.0 | 3.36 | - | - |
| **R00670.1** | Zaragizh | 1126 | ND | 9 | 75 | 2.38 | 0.21 | 11.3 | -20.4 | 20.7 | 68.5 | 3.9 | 4513 | 60 |
| **R00670.2** | Zaragizh | 1072 | ND | 9 | 90 | 9.32 | 0.87 | 11.0 | -19.7 | 18.5 | 53.3 | 3.4 | 4799 | 32 |
| **R00670.3** | Zaragizh | 1036 | ABA-UF | x | x | 22.48 | 2.17 | 10.8 | -19.6 | 15.1 | 42.8 | 3.3 | 4928 | 34 |
| **R00670.4** | Zaragizh | x | ND-UF | 9 | 90 | - | - | 10.6 | -19.7 | 16.2 | 42.9 | 3.1 | 4906 | 59 |
| **R00671.1** | Abri Cellier | 137 | ND | 9 | 75 | 0.29 | 0.21 | - | - | - | - | - | - | - |
| **R00671.2** | Abri Cellier | 136 | ND | 9 | 90 | 0.3 | 0.22 | - | - | - | - | - | - | - |
| **R00671.3** | Abri Cellier | 133 | ABA | x | x | 0.12 | 0.09 | - | - | - | - | - | - | - |

| Lab number | Site | Mass (mg) | P code | Extraction time (h) | Water temp °C | Collagen yield (mg) | % collagen yield | $\delta^{15}N$ (‰) | $\delta^{13}C$ (‰) | % N | % C | C:N atomic ratio | $^{14}C$ age (BP) | ± value (1 σ) |
|---|---|---|---|---|---|---|---|---|---|---|---|---|---|---|
| **R00672.1** | Abri Cellier | 351 | ND | 7 | 75 | 1.07 | 0.30 | - | - | - | - | - | - | - |
| **R00672.2** | Abri Cellier | 346 | ND | 7 | 90 | 1.15 | 0.33 | 8.2 | -20.5 | 11.5 | 35.4 | 3.6 | - | - |
| **R00672.3** | Abri Cellier | 342 | ABA-UF | x | x | 12.54 | 3.67 | 8.0 | -20.5 | 14.3 | 41.5 | 3.4 | - | - |
| **R00673.1** | Abri Cellier | 1536 | ND | 7 | 75 | 4.59 | 0.30 | 4.6 | -21.9 | - | - | 3.9 | 13528 | 44 |
| **R00673.2** | Abri Cellier | 1487 | ND | 7 | 90 | 5.56 | 0.37 | 5.0 | -20.2 | 13.5 | 40.1 | 3.5 | 22091 | 80 |
| **R00673.3** | Abri Cellier | 1391 | ABA-UF | x | x | 16.7 | 1.20 | 4.7 | -20.5 | 11.4 | 34.4 | 3.5 | 28251 | 190 |
| **R00677.1** | Denisova cave | 2388 | ND | 9 | 90 | 11.47 | 0.48 | 4.5 | -20.2 | 15.8 | 43.4 | 3.2 | - | - |
| **R00677.2** | Denisova cave | 2319 | ABA-UF | x | x | 0.17 | 0.01 | - | - | - | - | - | - | - |
| **R00678.1** | Denisova cave | 1712 | ND | 9 | 90 | 3.03 | 0.18 | 4.7 | -23.7 | 4.7 | 26.4 | 6.6 | - | - |
| **R00678.2** | Denisova cave | 1643 | ABA-UF | x | x | 0.32 | 0.02 | - | - | - | - | - | - | - |
| **R00679.1** | Denisova cave | 667 | ND | 9 | 90 | 24.35 | 3.65 | 7.8 | -18.9 | 15.6 | 42.7 | 3.2 | - | - |
| **R00679.2** | Denisova cave | 628 | ABA | x | x | 1.03 | 0.16 | 7.6 | -18.6 | 1.7 | 5.5 | 3.9 | - | - |
| **R00680.1** | Denisova cave | 914 | ND | 9 | 90 | 28.15 | 3.08 | 6.0 | -19.7 | 13.6 | 37.6 | 3.2 | > 47700 | - |
| **R00680.2** | Denisova cave | 877 | ABA-UF | x | x | 21.04 | 2.40 | 5.9 | -19.7 | 16.2 | 44.2 | 3.2 | > 48600 | - |
| **R00680.3** | Denisova cave | 914 | ND-UF | 9 | 90 | - | - | 5.3 | -19.7 | 17.8 | 48.0 | 3.1 | >42500 | |
| **R00681.1** | Denisova cave | 509 | ND | 9 | 90 | 3.25 | 0.64 | 9.7 | -17.9 | 13.3 | 37.2 | 3.3 | 30205 | 216 |
| **R00681.2** | Denisova cave | 497 | ABA-UF | x | x | 36.19 | 7.28 | 9.6 | -17.8 | 16.0 | 43.6 | 3.2 | 42949 | 1105 |

| Lab number | Site | Mass (mg) | P code | Extraction time (h) | Water temp °C | Collagen yield (mg) | % collagen yield | $\delta^{15}N$ (‰) | $\delta^{13}C$ (‰) | % N | % C | C:N atomic ratio | $^{14}C$ age (BP) | ± value (1 σ) |
|---|---|---|---|---|---|---|---|---|---|---|---|---|---|---|
| **R00682.1** | Denisova cave | 466 | ND | 9 | 90 | 8.93 | 1.92 | 6.5 | -19.6 | 16.0 | 45.1 | 3.3 | - | - |
| **R00682.2** | Denisova cave | 446 | ABA | x | x | x | 0.00 | - | - | - | - | - | - | - |
| **R00683.1** | Denisova cave | 515 | ND | 9 | 90 | 12.09 | 2.35 | 9.0 | -19.4 | 18.3 | 50.2 | 3.2 | 45301 | 1471 |
| **R00683.2** | Denisova cave | 490 | ABA-UF | x | x | 10.62 | 2.17 | 9.2 | -19.3 | 17.8 | 49.0 | 3.2 | >52100 | - |
| **R00683.3** | Denisova cave | 515 | ND-UF | 9 | 90 | - | - | 8.7 | -19.3 | 16.5 | 42.4 | 3.0 | > 38300 | - |

### 2. Methods

In this project, two collagen isolation methods were used, a non-destructive method using hot water extraction of whole bone, and the routine (destructive) method of demineralizing bone with dilute HCl. This approach enables comparison of the non-destructive and destructive methods.

#### 2.1. Routine (destructive) collagen extraction - HCl demineralization

As mentioned in the main text, sample sizes of archaeological samples were varied to simulate a realistic non-destructive sampling situation. This resulted in large differences in sample masses within batches. Depending on the bone's quality and composition, demineralization or gelatinization sometimes took longer, requiring duration adjustments or additional acid washes. Small samples or those with minimal material remaining after the ABA treatment did not continue to the ultrafiltration step and were freeze-dried directly after Ezee™ filtering at 90 µm. These samples are identified with the "ABA" pretreatment code, while the ultrafiltered, demineralized collagens are denoted "ABA-UF"(Table S1 and Table S2).

Normally, the low cut-off value for bone dating in the Higham lab is a collagen yield of ≤ 0.5% wt. However, to compare the destructive method with the non-destructive approach and assess the reliability of dates concerning material preservation, samples below this minimum were sometimes processed.

The first sample batch included specimens from Abri Cellier, Ipatovo and Zaragizh. Samples R00001.92 and R00001.93 are laboratory backgrounds. The same bone fragments that underwent consecutive hot water extractions were used for the destructive comparison. In this batch no additional acid washes were needed, and the protocol was followed as mentioned in the main text.

The second sample batch included samples from Denisova Cave. Bone fragments were cut in half, and one subsample was treated non-destructively while the other underwent the conventional, destructive approach. Samples R00001.103 and R00001.104 are the laboratory backgrounds. Again, samples showed great differences in behavior. After repeated acid washes in 0.5*M* HCl for demineralization, not every sample was completely demineralized. Sample R00678.2 underwent an additional acid wash the next day. Samples R00680.3 and R00677.2 underwent two additional acid washes.

In some cases, the ultrafiltration step was not applied in the destructive protocol, because of very low sample sizes or the fear of losing too much material. This applies to the following samples: R00001.92, R00667.3, R000671.3, R00679.2 and R00682.2. Information on precise pretreatment steps is also recorded in Table S2.

Apart from these examples, there were no deviations from the protocol.

### 2.2. Non-destructive collagen extraction --- hot water extraction of whole bone

#### 2.2.1. Experimental setup

For the hot water extraction, pre-baked Pyrex glassware (X-hours at 500°C) was used as well as a heating block (Corning® LSE™ Digital Dry Bath Heater) and hot plates (MR Hei-Standard by Heidolph Scientific Products GmbH). Smaller samples were loaded into 12mL borosilicate glass tubes, covered with MilliQ™ water and inserted into a heating block. Evaporation was reduced to a minimum by using glass lids.

Larger samples were placed in 30mL beakers on a hot plate. Beakers were covered with aluminium foil and a glass lens. It was difficult to completely reduce all evaporation from the beakers. It was therefore necessary to check the samples regularly and ensure that they were covered throughout the process. Additional MilliQ™ water was added if required.

In both cases the temperature was checked regularly with a thermometer to ensure uniform heating.

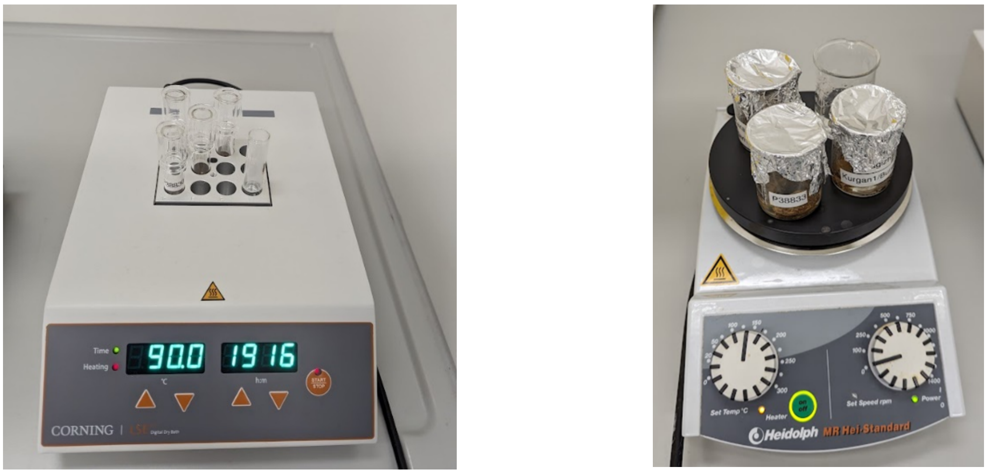

**Figure 1.** Experimental set up. Left, the heating block is used to treat small samples in 12mL tubes. The right picture shows bigger samples treated in 30mL beakers (photos by K. Luftensteiner).

#### 2.2.2. Detailed workflow and experimental set up changes

During the experimental programme, parameters such as temperature and extraction time were changed and certain steps were added e.g., Ezee filtering and ultrafiltration.

Initially, hot water extraction was done on 10 Hollis bones heated at 75°C for 1-to-10 hours. The bone fragments were treated in 12mL sample tubes. Three bone fragments whale bone were processed at 75°C for 2, 5 and 7 hours.

Based on these parameters, the method was then used on the first batch of archaeological samples. The batch comprised samples from Ipatovo 3 (*n*=1) and Zaragizh(*n*=3), both Bronze age sites in the north Caucasus (Russia) as well as Palaeolithic samples from Abri Cellier (*n*=3) (France). To simulate a realistic non-destructive sampling situation, where samples varied widely in shape and mass, we selected samples of varying size, from 133 to 2388 mg. Following the first heating experiment at 75°C we observed a lower than expected 'collagen' yield in several instances. We therefore increased the temperature to 90°C to test whether or not the yields could be increased (Figure S2 and Table S4). For this step the same bone fragments were used again. These results indicated that "collagen" yields were higher at 90°C. Compared to the Hollis bone these porous archaeological samples contained more sediment and particulate detritus and caused the water in the tubes to become cloudy. To remove these contaminants and clarify the solutions, the liquid was passed through 90 µm Ezee filters.

The second batch of archeological samples from Denisova Cave ($n$=7) (Russia) was treated non-destructively at 90°C. We divided the samples into two equal groups; one for the non-destructive treatment and a second for the routine destructive method. When collagen yields for the non-destructive method were high enough, ultrafiltration was used. Collagen from these samples was solubilised and then ultrafiltered. The application of the non-destructive protocol without ultrafiltration was denoted "ND" while those samples that underwent ultrafiltration are denoted "ND-UF". Starting weight, extraction yields, analytical data and radiocarbon ages are shown in Table S2.

### 2.3. Amino acid profiling

Samples of collagen for amino acid profiling were hydrolyzed in 6N HCl/1% phenol for 24 hrs at 110°C in vacuo, then dried under vacuum (Speed-Vac Labconco). Dried material was dissolved in Li Diluent (Pickering) containing 100nmol/mL AE-Cys (S-(2-Aminoethyl)-L-cysteine, Sigma A2636). 50ul of hydrolysate was loaded onto Hitachi 8080 Amino acid analyzer and subjected to strong cation exchange (Hitachi PF part #855-4515) to separate the amino acids. A secondary reaction with ninhydrin results in absorbance recorded at 570 nm and 440nm. A known standard run in the same sequence is used to create response factors to quantify each amino acid. Each sample contains AE-Cys (internal standard) to correct the results for any variation in sample volume (from the auto-sampler).

The modern collagen we are comparing the amino acid profiles of the Hollis standard with comes from a modern cow bone from 2010. It was degreased in warm water followed by chloroform-methanol extraction. The powder of the bone is used as a source of modern bone collagen. Since it is known that 20-50% wt % of collagen from archaeological bone is non-hydrolysable, one expects some variation in the proportions of amino acids measured in a profile such as this. This is because hydrolysis only releases amino acids from more intact parts of the bone skeleton with undegraded peptide bonds (see van Klinken, 1999). For this reason, amino acid profiles can sometimes be less than informative regarding the exact preservation of biomolecules that are derived from the bone or tooth under study and should be taken as a general guide (as we have done in this paper).

#### 2.4. Radiocarbon dating

Samples were combusted and graphitized using an Ionplus AGE3 graphitisation unit, pressed into aluminium targets and measured at the VERA (Vienna Environmental Research Accelerator) AMS facility, University of Vienna. (For details on aspects of target preparation and AMS measurement see Steier et al. [13] and Golser and Kutschera [14].)

Radiocarbon dates are reported in radiocarbon years BP (Before Present - AD 1950) using the Libby half-life of 5568 years. The results were normalized to typically seven separate graphitized samples of $^{14}C$ reference materials NIST Oxalic Acid II, IAEA-C3, IAEA-C6. No background correction was applied. The uncertainty given includes counting statistics, the reproducibility of repeated measurements on the samples, and the deviation of the reference materials from their nominal values. Isotopic fractionation in the AMS instrument has been corrected using the $\delta^{13}C$ value measured on the AMS. Carbon and nitrogen stable isotope and C/N values were measured independently using an EA-IRMS (Elemental Analyser/Isotope Ratio Mass Spectrometer, Thermo Scientific) to a precision of ± 0.3‰ relative to VPDB for carbon and ± 0.2‰ relative to AIR for nitrogen. Radiocarbon determinations from archaeological samples reported in this paper were given 'VIE' prefixes numbers denoting their completion.

#### 2.5. Programs and statistics

For calculation of the calibrated ages shown in Figure 3 and Figure S3 as well as the visualization of the plots the OxCal online program (version 4.4) was used[15,16] with the northern Hemisphere INTCAL dataset.

For statistical calculations and plotting Microsoft Excel and RStudio (version 2023.06.0) were used. The statistical comparisons of radiocarbon age determinations were calculated after Ward and Wilson (1978)[17].

### 3. Results

#### 3.1. Hollis Mammoth

Analytical data from the Hollis mammoth are documented in the main text and Table S2.

AMS results for the first application of hot water extracted material from the Hollis bone are shown in Table S3**Table S3**. No $^{14}C$ background correction was applied. The ages of destructively processed standards that were run in the same wheel are also presented. As mentioned in the main text, the results for the non-destructively processed standards are younger than expected when compared to the destructive treatment. The material dated has not been ultrafiltered or cleaned in any way, and therefore the $^{14}C$ measurements are not final.

**Table S3.:** Radiocarbon ages for Hollis bone samples analysed using the 75ºC hot water extraction and Destructive method approaches. Samples R00001.47 - R00001.58 and samples R00001.103 - R00001.69 were run in two consecutive AMS wheels due to their different time of treatment.

| Sample number | Chemistry | Extraction time, h | Sample name | $^{14}C$ age (RC yr. BP) | SD ± 1 σ |
|---|---|---|---|---|---|
| **R00001.47** | non-destructive | 1 | Hollis bone | 42282 | 176 |
| **R00001.48** | non-destructive | 2 | Hollis bone | 44006 | 183 |
| **R00001.49** | non-destructive | 3 | Hollis bone | 45204 | 213 |
| **R00001.50** | non-destructive | 4 | Hollis bone | 46034 | 241 |
| **R00001.51** | non-destructive | 5 | Hollis bone | 47250 | 332 |
| **R00001.52** | non-destructive | 6 | Hollis bone | 43895 | 188 |
| **R00001.53** | non-destructive | 7 | Hollis bone | 48068 | 249 |
| **R00001.56** | destructive | x | Hollis bone | 50241 | 285 |
| **R00001.57** | destructive | x | Hollis bone | 50031 | 273 |
| **R00001.58** | destructive | x | Hollis bone | 52313 | 377 |
| **R00001.103** | destructive | x | Hollis bone | 50626 | 497 |
| **R00001.104** | destructive | x | Hollis bone | 50971 | 544 |
| **R00001.67** | non-destructive | 8 | Hollis bone | 45584 | 517 |
| **R00001.68** | non-destructive | 9 | Hollis bone | 46398 | 328 |
| **R00001.69** | non-destructive | 10 | Hollis bone | 44991 | 308 |

#### 3.2. VIRI-I Whale bone

The second sample was VIRI-I whale bone, which had collagenous extracts with 0.82 to 3.22% yields (Table S2). Surprisingly, the lowest yield was obtained after 7 hours. The collagen was dark yellow, with a solid sticky consistency following freeze-drying. Analytical data on those fragments gave aberrant results with C/N ratios of 4.24 and 4.12 after 5 and 7 hours, respectively. The samples were considered 'failed' and were not radiocarbon dated.

#### 3.3. Archaeological samples

##### 3.3.1. Yields and analytical data

Two batches of archaeological samples were used to test the protocol. Samples rated as a 'fail' either had insufficient material for further analysis or the residue was poor quality and was yellow, sticky, and solid. Therefore, although some samples produced the same amount of material, not all were rated as successful and passed to the next stage.

The first batch of 7 samples from Zaragizh, Ipatovo and Abri Cellier and was treated between 7 to 9 hours at 75°C. Five of the 7 samples failed to produce any significant yield of 'collagen' (Figure S2a) and these were termed "failed" samples.

The temperature elevation from 75°C to 90°C resulted in significantly higher yields and a higher quality of collagen, based on C/N ratios, %C and other analytical proxies (see Figure S2b and c). Since the extraction process was carried out twice on the same bone fragment, the final reported yields include the yields of both extraction steps (Table S4). Following these results, we modified and adjusted our extraction protocol to the higher temperature (90°C) for all further testing.

Collagen yields of the archaeological samples were lower compared to the Hollis bone and is due to the better collagen preservation in Hollis. Sample R00670 from the Zaragizh site showed the highest non-destructive yield - a 1.04% wt. yield at 90°C. The Zaragizh sample was the only archaeological bone whose collagen yield exceeded the Higham lab's 0.5% minimum value for dating. However, < 0.5% wt. yield samples were processed for direct comparison with the destructive protocol. On average, at the extraction temperature of 90°C, the 7 samples yielded 0.54% collagen (*SD* = 0.26).

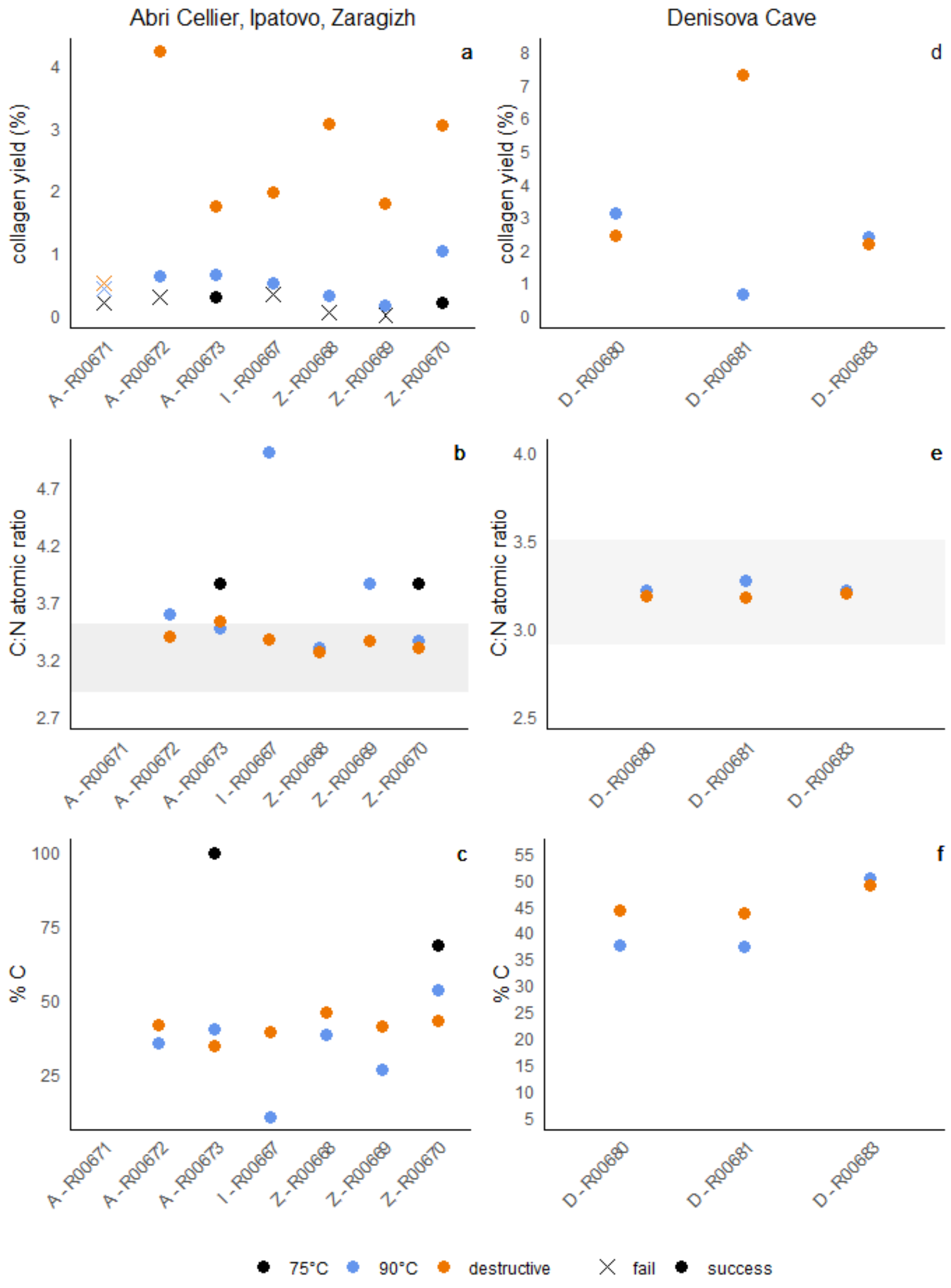

**Figure S2:** Analytical data from archaeological samples after non-destructive hot water extraction. The left column displays samples extracted at 75°C and 90°C (Abri Cellier, Ipatovo and Zaragizh), while the right column shows Denisova samples that underwent 90°C extraction only. None of the samples were ultrafiltered. From top to bottom, the rows represent collagen yield percentage (a and d), C/N ratios (b and e), and %C (c and f). Note that for Zaragizh, Itapovo and Abri Cellier results the extraction process was carried out twice on the same bone fragment, therefore the total reported yields of collagen include the yields of both extraction steps (Table S4). Sample R00668 is the human tooth.

Heating at 90ºC improved both collagen yields and C/N values. At 75ºC, samples R00670 (Zaragizh) and R00673 (Abri Cellier) had C/N ratios of 3.86 and 3.85, and C/N values of 3.36

and 3.46 with 90°C extraction. Our interpretation is the 90°C hot water extraction is mobilizing more, better preserved collagen than at 75°C.

For two samples (R00667 from Ipatovo and R00669 from Zaragizh), the 75°C fraction was too small for analysis, but the 90°C extracted collagen yielded very high C/N values of 5.0 and 3.85, respectively. The C/N values of destructive treatment residues were 3.37 and 3.36, and in the expected range.

The remaining samples (R00668, R00672, R00680) have C/N ratios, and %C and %N values, within the expected range when compared with the values for destructively extracted collagen (Table S2).

A second batch of archaeological samples comprised bones from the East Chamber at Denisova Cave, Russia (n=7). This location in the cave is known for moderate to good collagen preservation, with samples as old as 200 ka often yielding high-quality collagen. Each sample was split into two pieces; one-half underwent hot water extraction at 90°C, while the other half had its collagen extracted by demineralization in HCl. Three samples failed the destructive pretreatment, precluding direct comparisons; results from the non-destructive approach for those samples are shown in Table S2.

Three samples from Denisova cave yielded enough material to allow us to compare the non-destructive 90°C method (without ultrafiltration) against the routine destructive method (see Table S2 for more detail).

Samples R00680 and R00683 show similar results in yield, C/N ratio and %C. R00681 differs in collagen yield, where demineralization yielded significantly more collagen than by hot water extraction. We attribute this yield difference to the better preservation of the Denisova Cave samples.

**Table S4**: Collagen yields from the first archaeological sample batch used to test temperature effects. The upper part of the table shows the yields after each extraction isolated. Samples were weighed before each round of extraction. The lower part of the tables shows the total yield. Here the yields are documented under the assumption that collagen extracted from a previous treatment would also have gone into solution and was therefore added to the yield. For calculations of the %yield, the sample weight before the first extraction (75°C) was used.

**YIELDS**
**after each round of extraction**

| Sample | Site | 75°C bone mass (mg) | 75°C yield (mg) | 75°C yield (%) | 90°C bone mass (mg) | 90°C yield( mg) | 90°C yield (%) | Destructive bone mass (mg) | Destructive yield (mg) | Destructive yield (%) |
|---|---|---|---|---|---|---|---|---|---|---|
| R00668 | Zaragizh | 1229 | 0.56 | 0.05 | 1224 | 3.40 | 0.28 | 564 | 33.70 | 5.98 |
| R00669 | Zaragizh | 1077 | 0.08 | 0.01 | 1051 | 1.79 | 0.17 | 1046 | 17.28 | 1.65 |
| R00670 | Zaragizh | 1126 | 2.38 | 0.21 | 1072 | 9.32 | 0.87 | 1036 | 22.48 | 2.17 |
| R00667 | Ipatovo | 191 | 0.64 | 0.34 | 187 | 0.34 | 0.18 | 187 | 2.77 | 1.47 |
| R00671 | Abri Cellier | 137 | 0.29 | 0.21 | 136 | 0.30 | 0.22 | 133 | 0.12 | 0.09 |
| R00672 | Abri Cellier | 351 | 1.07 | 0.30 | 346 | 1.15 | 0.33 | 342 | 12.54 | 3.67 |
| R00673 | Abri Cellier | 1536 | 4.59 | 0.30 | 1487 | 5.56 | 0.37 | 1391 | 16.70 | 1.2 |

**TOTAL YIELDS**
**All values added**

| Sample | Site | Starting weight (mg) | 75°C yield (mg) | 75°C yield (%) | 75°C+90°C yield (mg) | 75°C+90°C yield (%) | 75°C+90°C+destructive yield (mg) | 75°C+90°C+destructive yield (%) |
|---|---|---|---|---|---|---|---|---|
| R00668 | Zaragizh | 1229 | 0.56 | 0.05 | 3.96 | 0.32 | 37.66 | 3.06 |
| R00669 | Zaragizh | 1077 | 0.08 | 0.01 | 1.87 | 0.17 | 19.15 | 1.78 |
| R00670 | Zaragizh | 1126 | 2.38 | 0.21 | 11.70 | 1.04 | 34.18 | 3.04 |
| R00667 | Ipatovo | 191 | 0.64 | 0.34 | 0.98 | 0.51 | 3.75 | 1.96 |
| R00671 | Abri Cellier | 137 | 0.29 | 0.21 | 0.59 | 0.43 | 0.71 | 0.52 |
| R00672 | Abri Cellier | 351 | 1.07 | 0.30 | 2.22 | 0.63 | 14.76 | 4.21 |

| | | | | | | | | |
|---|---|---|---|---|---|---|---|---|
| R00673 | Abri Cellier | 1536 | 4.59 | 0.30 | 10.15 | 0.66 | 26.85 | 1.75 |

#### 3.3.2. AMS results on archaeological samples

Both destructive and non-destructively extracted collagen samples were AMS dated (Figure S3 and Table S5). The data was corrected for $^{14}C$ laboratory background by subtracting the Fm result of the corresponding Hollis bone samples. The Zaragizh samples agree with the expected Bronze Age estimate. Some age differences were apparent between the non-destructively and destructively processed samples. Sample R00681 (Denisova Cave) produced statistically different ages ($T = 128.11$; df = 1, $\chi^2 = 3.84$) 42,949 ± 1105 BP (VIE-1169) by destructive demineralization and 30,205 ± 216 BP (VIE-1168) by hot water extraction. Sample R00673 (Abri Cellier), was hot-water-extracted twice, at 75°C and 90°C. Collagen extracted at 75°C dated 13,528 ± 44 BP (VIE-1162), while the 90°C-extracted collagen dated to 22,091 ± 80 BP (VIE-1163). Collagen extraction by HCl demineralization was significantly older, 28,251 ± 190 BP (VIE- 1164). Both the 75°C and 90°C non-destructive results are statistically different from the destructive approach ($T = 5698.99$; df=1, $\chi^2 = 3.84$ ; $T$=892.84; df=1;$\chi^2$=3.84, respectively). This underscores the importance of using post-extraction purification steps such as ultrafiltration or XAD-2 resin treatment, that removes contaminants and results in accurate $^{14}C$ measurements.

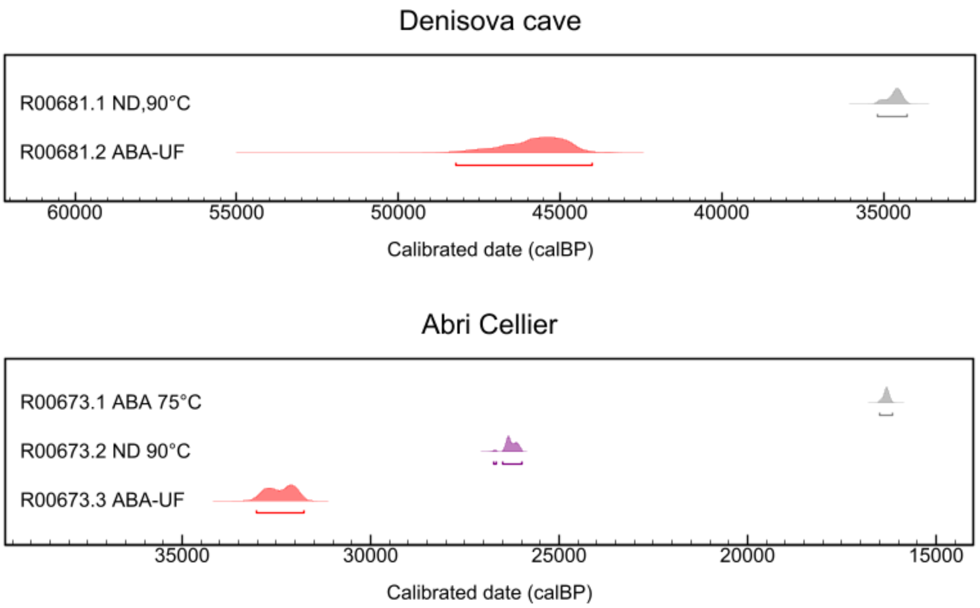

**Figure S3:** Calibrated ages (cal BP) of the samples that were radiocarbon dated. 'ND' denotes the non-destructive treatment, 'ABA' for the conventional, destructive approach. '-UF' marks samples that were post-purified using ultrafiltration.

**Table 5 :** Pretreatment information and radiocarbon ages for bones that were not ultrafiltered after the non-destructive hot water extraction. P code denotes the pretreatment method. The code 'ND' stands for the non-destructive treatment, while 'ABA-UF' denotes samples that were treated destructively and were ultrafiltered. A direct age comparison for these samples is therefore not possible.

| R_number | site | P code | Time (h) | °C | VIE ID | $^{14}C$ age BP | ± 1 σ |
|---|---|---|---|---|---|---|---|
| R00673.1 | Abri Cellier | ND | 7 | 75 | VIE-1162 | 13528 | 44 |
| R00673.2 | Abri Cellier | ND | 7 | 90 | VIE-1163 | 22091 | 80 |
| R00673.3 | Abri Cellier | ABA-UF | x | x | VIE- 1164 | 28251 | 190 |
| R00681.1 | Denisova cave | ND | 9 | 90 | VIE-1168 | 30205 | 216 |
| R00681.2 | Denisova cave | ABA-UF | x | x | VIE-1169 | 42949 | 1105 |

#### 3.4. Morphological changes

Macroscopically, no morphological or dimensional changes were evident after hot water extraction was performed on the archaeological samples (Figure S4).

#### 3.5. Failed samples

As mentioned earlier, the interlaboratory standard, VIRI-I, showed aberrant results on all analytical data (Table S2). For this reason, the samples were considered as failed and were not radiocarbon dated.

The results of the destructive treatment for Denisova samples were poor and 3 samples failed to yield enough measurable collagen. Surprisingly, the non-destructive method did produce enough collagen by comparison. As mentioned previously, sample sizes differed greatly in this batch and therefore demineralization times ranged accordingly. We suspect these three samples did not contain collagen because they had variable biomolecular preservation, their processing times were too short or too long a combination of these factors

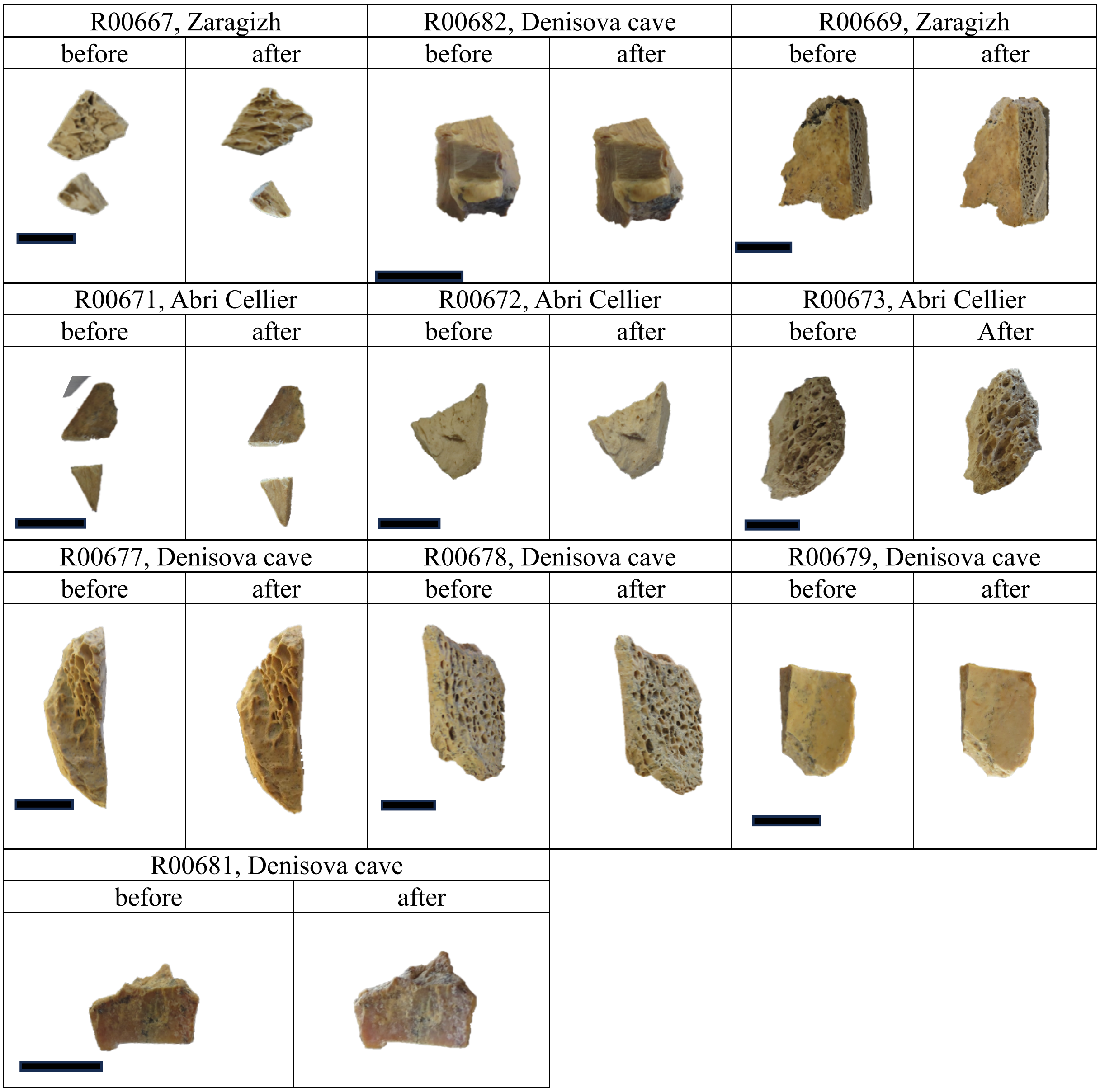

**Figure S4:** Pictures of bones before and after hot water extraction. Note that pictures were taken at slightly different angles and light settings. For the tooth sample, R006698, only the root was covered with MilliQ™ water, the crown was solely influenced by evaporation. The black bar equals 1cm.

### 4. Discussion

In the Destructive method, collagen yields $\leq$1.0-0.5% wt. are considered too low for accurate radiocarbon dating due to the effects of contamination[18]. Dobberstein et al.[19] reached similar conclusions, where minimal changes in quality control parameters were observed until collagen loss reached 99%. In our study, we applied the non-destructive protocol to samples with collagen yields within or below this range, to test whether these thresholds are applicable to novel extraction methods.

All non-destructively treated samples that yielded enough collagen for analytical data were near or below the cut off value of ~0.5% wt. The destructive yields, however, were not below that range. However, for some problematic samples where the total collagen yield (summed across multiple extraction rounds) exceeded this threshold, the abnormal quality control values may be because of lower extraction temperatures rather than insufficient collagen yield. This shows that the % collagen yield in case of non-destructive treatment is not an indicator for the overall preservation of a sample and underscores the importance of further refining the temperature and yield thresholds for reliable collagen extraction in low-collagen samples.

High C/N ratios were measured on hot water extracts that had not been purified by ultrafiltration. We think that high molecular weight collagen fragments can be mobilized more efficiently by an only acid-based treatment as these peptides are more securely bound within the bone matrix itself. Regardless, our data show that more collagen could be mobilized by increasing the treatment temperature from 75ºC to 90ºC.

Non-destructive sampling techniques for teeth are already known in the field of ancient DNA. In a study testing a minimally destructive protocol for ancient teeth[20] only the root was covered by the extraction buffer. This method produced ancient DNA comparable quality to samples treated destructively. However, the samples showed slight changes in degradation after the treatment, because diluted bleach and UV light irradiation was used for cleaning.

Another study focused on thermal degradation of DNA in human teeth[21]. Human teeth were exposed to temperatures of 100, 200 and 400°C for 60 minutes. Using Real-Time qPCR, DNA was detected in 100% of samples treated at 100°C, 80% in 200°C and 20% at 400%. However, most of the DNA was already degraded at temperatures between 100 and 200°C. Although no ancient samples were used in the DNA study, the results are an indication that dental material can tolerate very high temperatures. To evaluate the influence of our hot water treatment on aDNA in ancient teeth, further research is needed.

However, we show, albeit in one example, that when teeth are well-preserved, sufficient collagen can be extracted non-destructively from the root of human teeth and accurately dated even without ultrafiltration or other purification steps.
